# Zwitterionic polymer coating enabled chronic dopamine sensing and electrophysiology recording in free-moving mice

**DOI:** 10.64898/2026.02.08.704618

**Authors:** Bingchen Wu, Cort Thompson, Thomas Deakin, Yan Xu, Colleen McClung, Xinyan Tracy Cui

## Abstract

The brain’s complex network relies on both electrical and chemical signaling to support its physiological and cognitive functions. To fully understand neural circuit dynamics and their dysfunctions, it is crucial to simultaneously detect neurotransmitters and modulators alongside electrophysiological signals. The striatal dopamine circuits are integral to neurological processes such as movement, reward, learning, and circadian rhythm regulation, making it highly desirable to monitor both neural activity and dopamine (DA) levels in freely behaving animals. One promising approach involves the implantation of multimodal microelectrode arrays (MEAs). However, chronic electrochemical sensing of DA in freely moving animals faces significant challenges, including biofouling of sensing electrodes and the instability of Ag/AgCl reference electrodes. In this study, we developed two complementary strategies—surface grafting and photo crosslinking—to coat the MEA and implanted Ag/AgCl reference electrodes, respectively, with zwitterionic poly(sulfobetaine methacrylate) (PSB). The surface-grafted thin PSB coating effectively inhibits protein fouling and inflammatory responses to the MEA, while the PSB hydrogel protects the Ag/AgCl electrodes from delamination in vivo, ensuring a stable reference potential. By coating both the Ag/AgCl reference electrodes and flexible polyimide MEAs with PSB and PEDOT/CNT, we achieved stable DA detection and electrophysiological recordings in freely moving mice over a four-week period. Weekly electrochemical impedance spectroscopy confirmed the long-term stability of the implanted electrodes. Our method enables multidimensional analysis of behavioral patterns, electrophysiological activity, and DA dynamics, providing a comprehensive approach for neuroscience research. This work advances neurochemical and electrophysiological methodologies by offering reliable tools for longitudinal investigations of brain function in freely behaving animals.

## Introduction

The brain is the most complex organ in the human body, comprising tens of billions of neurons interconnected by hundreds of trillions of synapses. Information processing in neural circuits relies on two fundamental signaling modalities: electrical signals, such as action potentials, and chemical signals, including neurotransmitters and neuromodulators. The coordinated spatiotemporal interplay between these electrical and chemical signals underlies physiological processes and higher-order cognitive functions such as movement, learning, and memory[1-7]. Dysregulation of neurochemical signaling has been implicated in a wide range of neurological and neuropsychiatric disorders[8-12]. Given that hundreds of neurochemicals contribute to normal brain function, there is a critical need for technologies capable of monitoring neurochemical dynamics in conjunction with electrophysiological activity to better understand neural circuit function under both physiological and pathological conditions.

Among neuromodulatory systems, dopaminergic circuits—particularly those within the striatum—play a central role in regulating movement, reward processing, learning, and motivation [2-6, 13-15]. Dopamine (DA) signaling is also known to interact with broader brain-state variables, including arousal, sleep–wake transitions, and behavioral context [7, 16-19]. These functions are mediated by DA dynamics that unfold over multiple timescales and are tightly coupled to neural activity patterns. Consequently, simultaneous measurement of DA levels and electrophysiological signals in freely moving animals is essential for elucidating how neuromodulatory signals shape circuit-level computations [20, 21]. The *Clock*Δ19 mice model is known to have hyperactive manic behaviors, and hyper-dopamine activities, making them an ideal model to validate the multi-modal microelectrode arrays (MEAs) recording capability. Flexible MEAs coated with conducting polymers such as poly(3,4-ethylenedioxythiophene) doped with acid-functionalized carbon nanotubes (PEDOT/CNT) have emerged as promising platforms for multimodal neural interfacing. We have demonstrated that such devices can support stable dopamine detection using square-wave voltammetry (SWV) while simultaneously enabling electrophysiological recordings over extended periods. Using this technique, we have shown how the *Clock*Δ19 mutation, a mutation in the core circadian gene *Clock* which leads to functional knock-out of function, affects striatal DA levels [21]. These prior demonstrations, however, were primarily conducted in anesthetized animals [22]. Anesthesia is known to profoundly alter brain-wide electrophysiological activity, neuromodulator release, and network dynamics [23-27]. Moreover, both neural activity and neurotransmitter signaling differ substantially between anesthetized, sleeping, and awake states [28-31]. As a result, extending multimodal sensing capabilities to awake, freely moving animals is critical for eliminating anesthetic confounds and for linking neural and neurochemical dynamics to naturalistic behaviors.

Continuous and prolonged electrophysiological recordings electrophysiological recordings in freely moving animals are now well established, with extensive literature addressing surgical approaches, device integration, and mitigation of motion-related artifacts[32-36]. In contrast, chronic electrochemical sensing of tonic dopamine in freely moving animals remains far less developed [37, 38]. A major technical bottleneck lies in the long-term stability of the reference electrode. Ag/AgCl electrodes are widely used as reference electrodes due to their well-defined redox equilibrium and stable potential that is largely insensitive to current polarization and pH fluctuations. However, maintaining a stable AgCl layer in vivo over extended periods is challenging. Biofouling, de-chlorination, and mechanical degradation can cause implanted Ag/AgCl electrodes to deviate from ideal nonpolarizable behavior, leading to signal drift and degraded measurement fidelity [39-41]. To mitigate these issues, various protective coatings— such as Nafion, polyurethane, and cell membrane–derived layers—have been explored to extend Ag/AgCl electrode lifetimes in vivo [42-44]. Each approach, however, presents limitations. Nafion’s anion-exclusion properties may disrupt chloride ion equilibrium[45], polyurethane lacks robust antifouling performance, and biologically derived coatings may provoke immune responses.

In our previous work, we addressed reference electrode instability by using a removable Ag/AgCl wire inserted subcutaneously prior to each measurement [21, 41]. While effective for anesthetized experiments, this approach is impractical for chronic sensing in awake and freely behaving animals. Achieving reliable dopamine sensing in this context requires an implanted reference electrode that is mechanically stable, resistant to biofouling, and compatible with integrated head-mounted electronics.

Zwitterionic polymer coatings offer a promising solution to these challenges. Zwitterionic polymers contain equal numbers of positively and negatively charged moieties within each monomer unit, resulting in an overall neutral charge while maintaining a high local ionic density. This unique structure confers exceptional hydration and ultra-low-fouling properties. Zwitterionic materials have been widely applied in drug delivery [46], smart materials [47], lubrication coatings [48-50], and biosensors [51-53]. Strongly bound hydration layers formed at the polymer surface act as a physical and energetic barrier to biofouling [54-62]. Our previous studies have demonstrated that zwitterionic polymers can prevent non-specific protein adsorption, enzyme interaction, and cell adhesion both in vitro and in vivo, and reduce acute inflammatory brain tissue response to implants [59, 63, 64].

Building on this foundation, the present work evaluates whether zwitterionic poly(sulfobetaine methacrylate) (PSB) coatings can enable stable, long-term dopamine sensing and electrophysiological recording in freely moving animals [22]. We employ two complementary PSB-based strategies: a photo-crosslinked PSB hydrogel coating on Ag/AgCl reference electrodes to prevent delamination and biofouling, and a thin, surface-grafted PSB layer on flexible polyimide MEAs using a previously reported photoiniferter-based approach [63, 65]. PEDOT/CNT is subsequently deposited onto the electrode sites to enable high-quality electrochemical and electrophysiological measurements. Both PSB coated Ag/AgCl reference electrode and flexible MEA were implanted in the *Clock*Δ19 mutant mouse brain for 4 weeks. Electrochemical impedance spectroscopy (EIS) was measured weekly with both a fresh subcutaneously inserted Ag/AgCl and the implanted Ag/AgCl reference electrode to monitor the stability of the implanted reference electrode. Electrophysiology data were collected both with mice under anesthesia and freely moving. The recording quality was compared between non-coated MEAs from previous work [21] and the PSB coated MEAs with anesthetized electrophysiology recording. DA sensing was performed weekly with the animal freely moving in an open field box. At the 4-week time point, the animal received an I.P cocaine injection, and an additional 30 mins DA sensing was performed. During all free-moving experiments, animal movements were video recorded and offline tracked and processed for behavior measurements. This setup was tested in two proof-of-concept experiments to demonstrate the potential applications in circadian rhythm and pain studies. Collectively, this work establishes a robust materials-based strategy for chronic multimodal neural interfacing in freely behaving animals, advancing experimental capabilities for systems neuroscience research.

## Results and Discussion

### Deposition and Characterization of PSB Coating on Ag/AgCl Reference Electrodes and Polyimide MEAs

Since the surface chemistry of Ag/AgCl wire and polyimide is drastically different. Two different coating methods were utilized (Figure 1). For Ag/AgCl electrode, a dip coating strategy was used to coat PSB on the Ag/AgCl wires. A solution containing partially crosslinked PSB, dithiothreitol (DTT) crosslinker, and α-ketoglutaric acid photo-initiators was first prepared, then freshly made Ag/AgCl wires were manually dipped into the solution and quickly pulled out, exposed to UV lights for 30s, then repeated the process for 3 times to reach micron thickness coatings that can be visualized under SEM (Figure 1, a).

**Figure 1.**
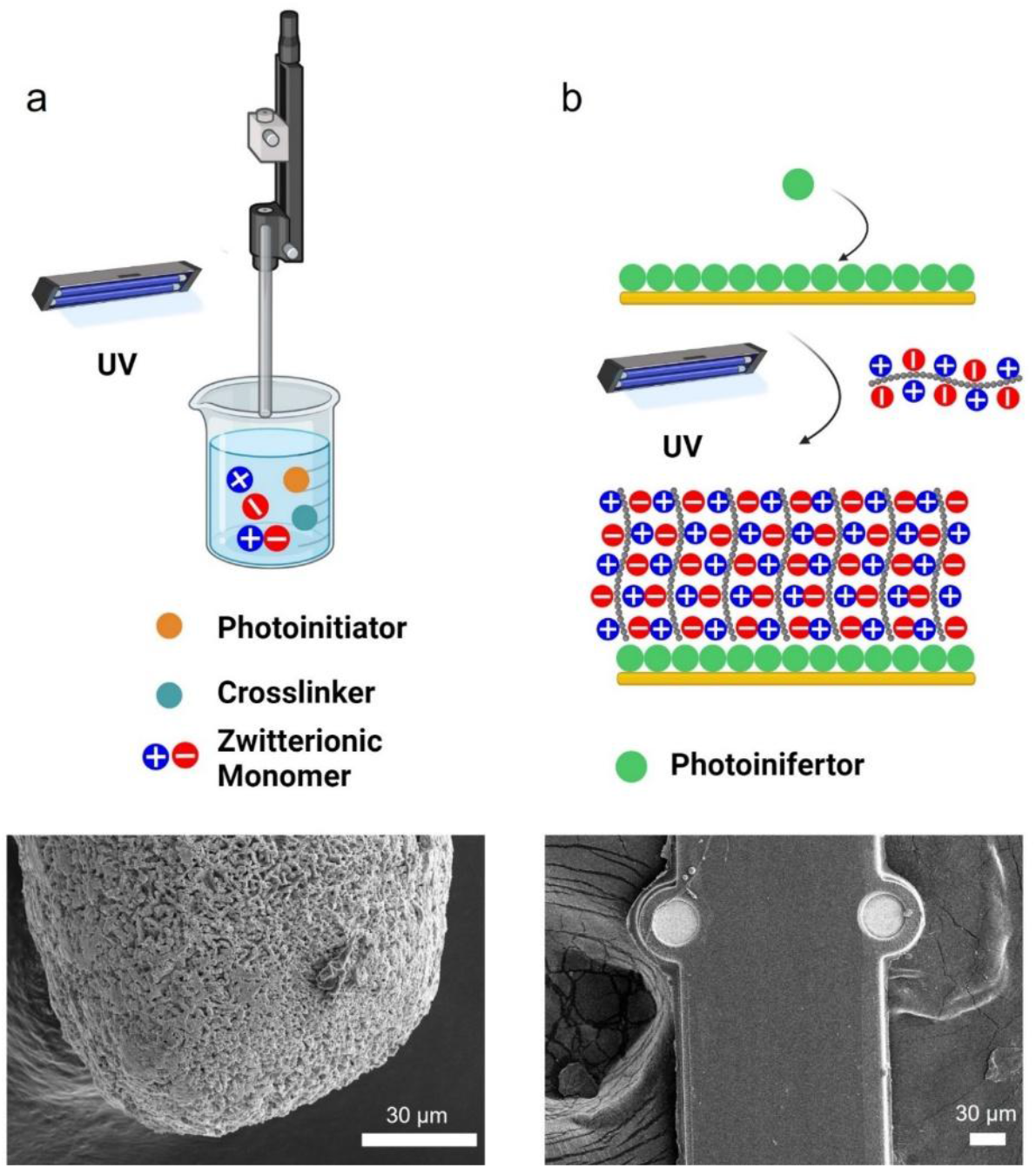
Schematic demonstration of PSB coating process and SEM of coated substrate. (a), Ag/AgCl were first dip into solution containing partially crosslinked PSB, photoinitiator α-ketoglutaric acid, and crosslinker DTT, then pulled out and exposed to UV lights. The process is repeated until desired thickness is reached. The SEM of PSB coated Ag/AgCl wire showed an organic polymer layer over the AgCl crystal structure. (b), The polyimide surface was first treated with O2 plasma, then a self-assemble-monolayer N,N-(Diethylamino) dithiocarbamoyl-Benzyl (trimethoxy silane) (SBDC) photoinifertor was immobilized on the surface. The SBDC-coated substrates were immersed in a monomer solution of PSB and irradiated with UV. This method produces tens of nanometer-thick PSB films uniformly of the MEA shank that is challenging to visualize under SEM.[22]

For coating polyimide MEAs, we adapted a surface-initiated polymerization strategy from previous work [63, 65] . The polyimide surface was first treated with O_2_ plasma, then a self-assemble-monolayer (SAM) N,N-(Diethylamino) dithiocarbamoyl-Benzyl (trimethoxy silane) (SBDC) photoinifertor was immobilized on the surface. Then the SBDC-coated substrates were immersed into a monomer solution of SBMA and irradiated with UV to start the surface-initiated polymerization process. This method produces tens of nanometer-thick PSB films uniformly on the MEA shank that is challenging to visualize under SEM (Figure 1, b). Thus, additional Fourier transform infrared spectroscopy (FTIR) and water contact angle (WCA) measurements were used to help validate whether the PSB was successfully grown on the polyimide surface (Figure 2). FTIR shows characteristic peaks of S=O from the sulfonate functional group of PSB (Figure 2, a), and WCA from PSB coated surface is significantly lower than non-coated polyimide, indicating a much more hydrophilic surface (Figure 2, b). Both results indicate PSB coatings have been successfully grown onto the polyimide surface.

**Figure 2.**
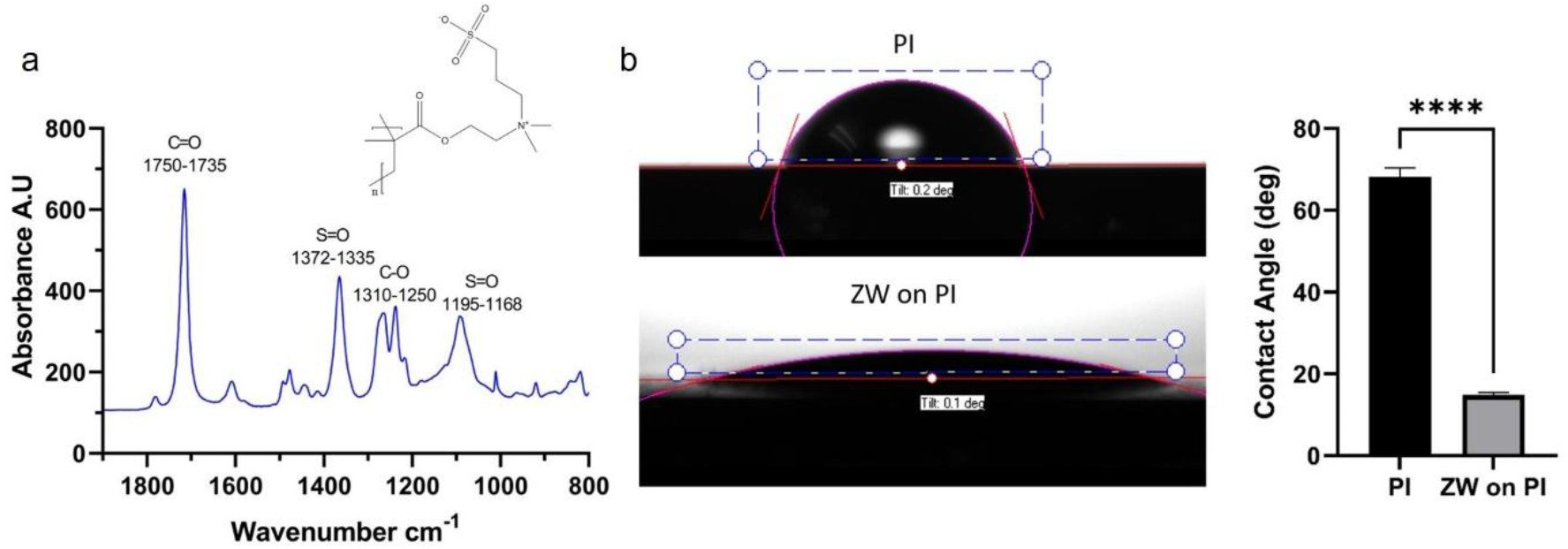
FTIR and WCA measurements of PSB-coated polyimide surface. (a), Polyimide(PI) spin coated on Si wafer. Cut into 3×5 cm rectangle pieces. FTIR was collected with PI coated Si wafer as background. FTIR shows characteristic peaks of C=O and C-O bonds from the ester functional groups. S=O from the sulfonate functional group of PSB. (b), The PI surface has a WCA of 68.18°±2.16°, after coating PSB on the PI surface, the WCA significantly decreased to 14.83°± 0.58°. n=3, mean±sem. Welch-t test. **** p<0.0001.[22]

The EIS of PSB-coated MEAs was measured and compared to pristine MEAs (Figure 3, a). The PSB coating did increase the EIS of the Pt sites, which could be attributed to a small amount of PSB being grown on the Pt surface. The SBDC SAM layer immobilization was achieved through a silanization reaction that can also happen on the Pt surface to a small degree [66, 67]. However, after the plasma treatment, the polyimide surface will contain much richer content of oxygen-containing functional groups such as hydroxy and carboxyl groups that are crucial silanization reactions. Thus, the polymer will be mostly coated on the polyimide surface. Also, after PEDOT/CNT were coated on the Pt sites, the PSB-coated MEAs had lower impedance than the PEDOT/CNT coated MEAs without PSB, indicating the PSB on Pt sites might have adhesion promotion effects for PEDOT/CNT coatings (Figure 3, a). SEM comparison of a PSB-coated MEA (Figure 3, b), a PEDOT/CNT coated MEA (Figure 3, c), and a PSB and PEDOT/CNT coated MEA (Figure 3, c) is shown in Figure 3. A clear feature of PSB-coated MEA is the smoother insulation surface (Figure 3, b & d). The PEDOT/CNT coated sites appeared darker than bare Pt sites (Figure 3 b vs Figure 3, c-d). This is because PEDOT/CNT is intrinsically less conductive than pure metal.

**Figure 3.**
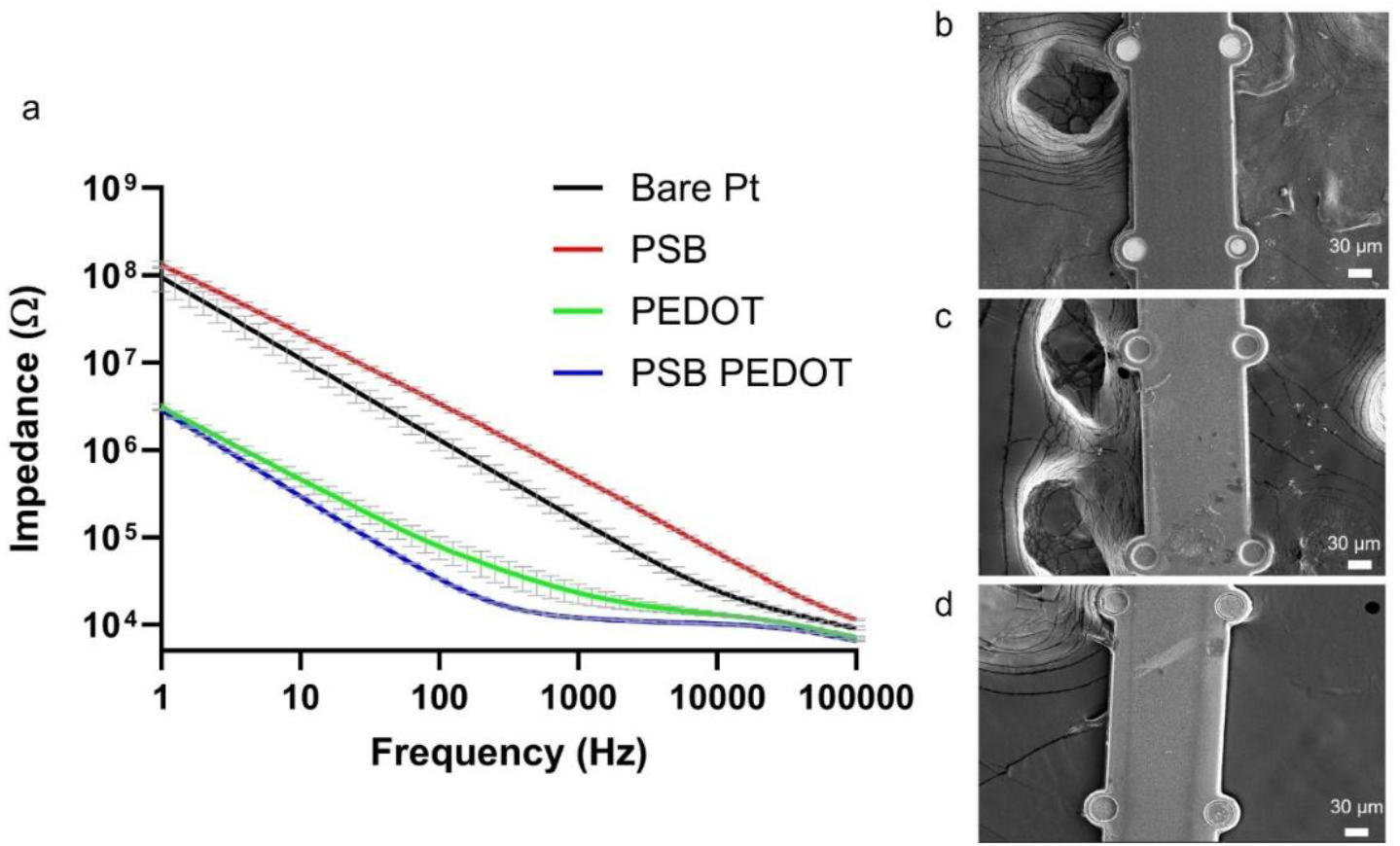
EIS and SEM characterization of PI MEA in different coating conditions. (a), EIS measurements of pristine MEA, PSB coated MEA, PEDOT/CNT coated MEA, and PSB+PEDOT/CNT coated MEA. (b), SEM of PSB-coated MEA. (c), SEM of PEDOT/CNT coated MEA. (d), SEM of PSB+PEDOT/CNT coated MEA.[22]

### Electrochemical Performance of PSB-coated Ag/AgCl and PSB PEDOT/CNT Coated MEA

One of the concerns about using PSB coatings on Ag/AgCl is its impact on ion transfers that could cause kinetic changes in electrochemical measurements. To test its electrochemical performance, we performed cyclic voltammetry (CV) scan of different scan rates in a 10mM ruthenium hexamine Ru(NH_3_)_6_^3+^ solution with a glassy carbon working electrode, a Pt counter electrode, and an Ag/AgCl reference electrode with or without PSB coatings (Figure 4 a). At both 1 v/s high scan rate and 0.1v/s low scan rate, the PSB-coated Ag/AgCl reference had a very slight potent shit to the cathodic side, indicating the coating might have effects of PSB coating on the ion equilibrium state that shifted the Ag/AgCl potential [45]. Importantly, the peak separation was maintained, indicating the electron transfer kinetics were not affected which is important for in vivo SWV DA sensing applications (Figure 4, b). Representative SWV calibration of PSB PECOT/CNT coated MEAs is shown in Figure 4, c. Robust response towards a gradient of physiologically relevant DA concentration is observed (Figure 4, c). Next, the SWV DA sensing of PSB PEDOT/CNT coated MEAs was compared with PEDOT/CNT coated MEAs (Figure 4 d). The PSB PEDOT/CNT coated MEA showed similar sensitivity compared to PEDOT/CNT coated MEA, indicating the PSB coating doesn’t affect the sensors’ sensitivity towards DA (Figure 4 d).

**Figure 4.**
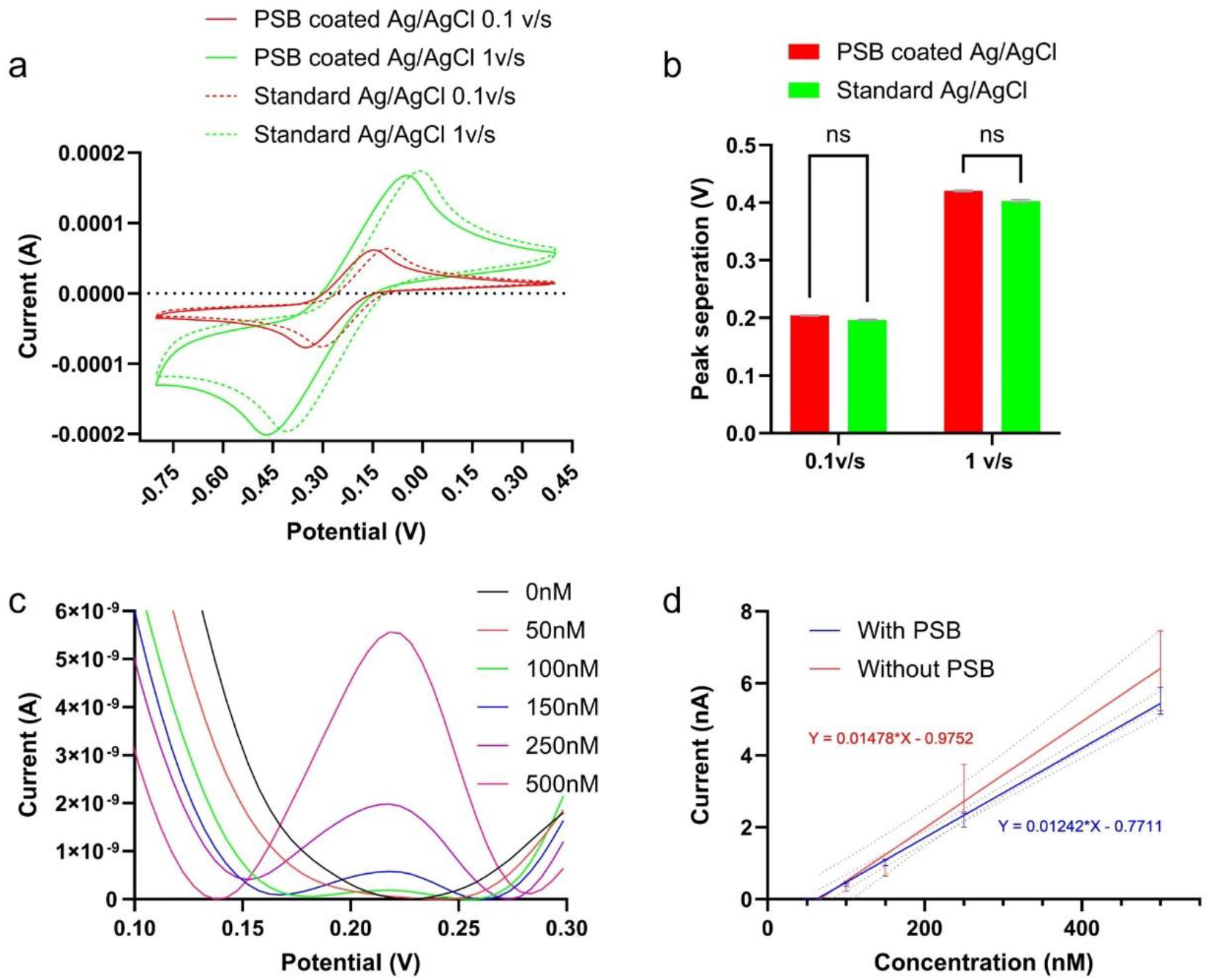
In vitro characterization of PSB coated Ag/AgCl referece electrodes and PSB PEDOT/CNT coated MEA. (a), CV scan of different scan rates in a 10mM Ru(NH3)63+ solution. A glassy carbon electrode is used as working, a Pt wire is used as counter, and an Ag/AgCl wire with or without PSB is used as reference. (b), Peak separation quantification for PSB coated and non-coated Ag/AgCl. Multiple t-test, Bonferroni posthoc, n =3. (c), Average SWV waveform of PSB PEDOT/CNT coated MEA during a DA calibration test (n=20). (d), Calibration curve comparison between PSB PEDOT/CNT and PEDOT/CNT only MEA. The sensitivity is calculated as the slope of the calibration curve. (n=20 for both groups). [22]

### *In Vivo* Stability of PSB PEDOT/CNT Coated MEA and PSB Coated Ag/AgCl

The surgical procedure was very similar to the previous study, except an additional cranial window was opened for PSB-coated Ag/AgCl reference implantation (Figure 5, a)[21]. For each weekly session, the animal was first anesthetized with isoflurane, and then EIS measurements were taken using both a fresh subcutaneous Ag/AgCl and the implanted PSB-coated Ag/AgCl reference. Electrophysiology data were also collected to directly compare recording quality with previous non-coated MEA [21]. The subcutaneous Ag/AgCl was removed before transferring animals into the open field box. Animals are allowed to fully recover from anesthesia for 30 mins (Figure 5, b). Then the SWV scanning started at the same time as the video recording which lasts 20 mins weekly. After finishing free moving DA sensing, the video recording was stopped and the headstage for the potentiostat connection was disconnected from the system. Electrophysiology recording was then started again, and the time stamp of the start of the video recording was synced to the recording system through a TTL trigger. Only on week 4, after the 5 mins free-moving electrophysiology session, the animal was gently taken out of the box and injected with cocaine. The animal was set free in the open field again and 30 mins of SWV free moving DA sensing was performed. Animals were sacrificed at the end of week 4.

**Figure 5.**
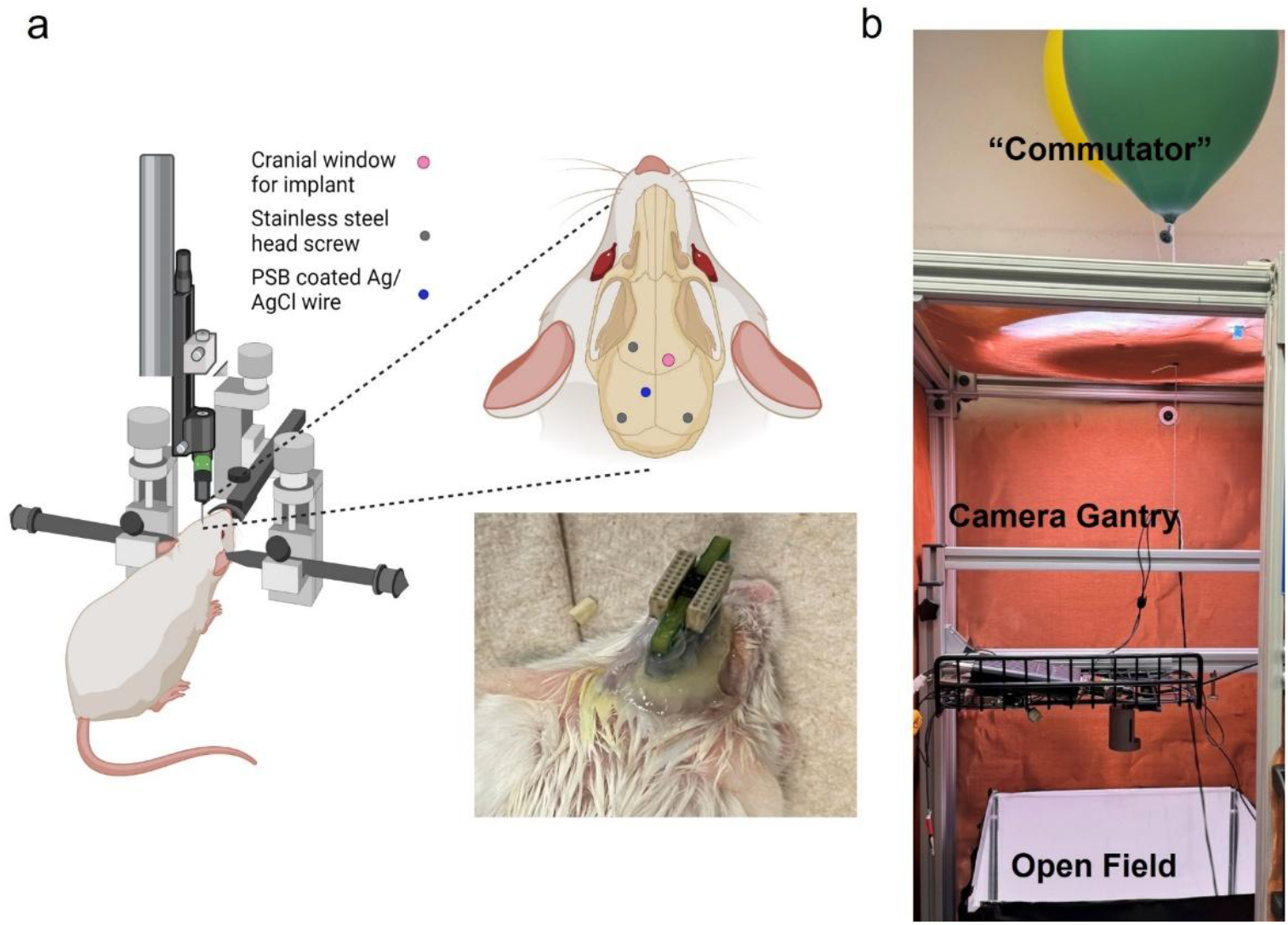
Surgical set-up and free-moving set-up. (a). Mice were anesthetized with isofluorane and placed under a stereotaxic frame. Three head screws were used as anchoring points for the dental cement fixture, and two of them on the contralateral side of the implant were used as ground for electrophysiology and electrochemistry measurement respectively. PSB-coated Ag/AgCl was implanted manually with forceps. A customized PCB with 2 omnetics connectors was used to connect MEAs with the recording and potentiostat system, respectively. b). Free-moving set-up. The cables for head-stage connections were levitated with helium balloons to avoid tangling/dragging during animal movements. A gantry is positioned in the middle of the open filed box for a center camera position. The whole set-up is enclosed inside a faraday cage. [22]

The impedance measured with subcutaneous Ag/AgCl and the implanted PSB Ag/AgCl was compared to gauge the electrochemical stability of the implant reference. The measured impedance difference between the two groups was relatively larger at Day 0 and Day 7 possibly due to additional inflammation response surrounding the PSB Ag/AgCl implantation site.

Although coated with antifouling coating, the large diameter of the Ag/AgCl wire can cause extensive trauma during initial surgery. Similar impedance measurements were observed at later time points between the two groups as the initial injury response cleared away, indicating PSB coating can protect Ag/AgCl reference from biofouling and ion contamination (Figure 6, a) [39-41, 45].

**Figure 6.**
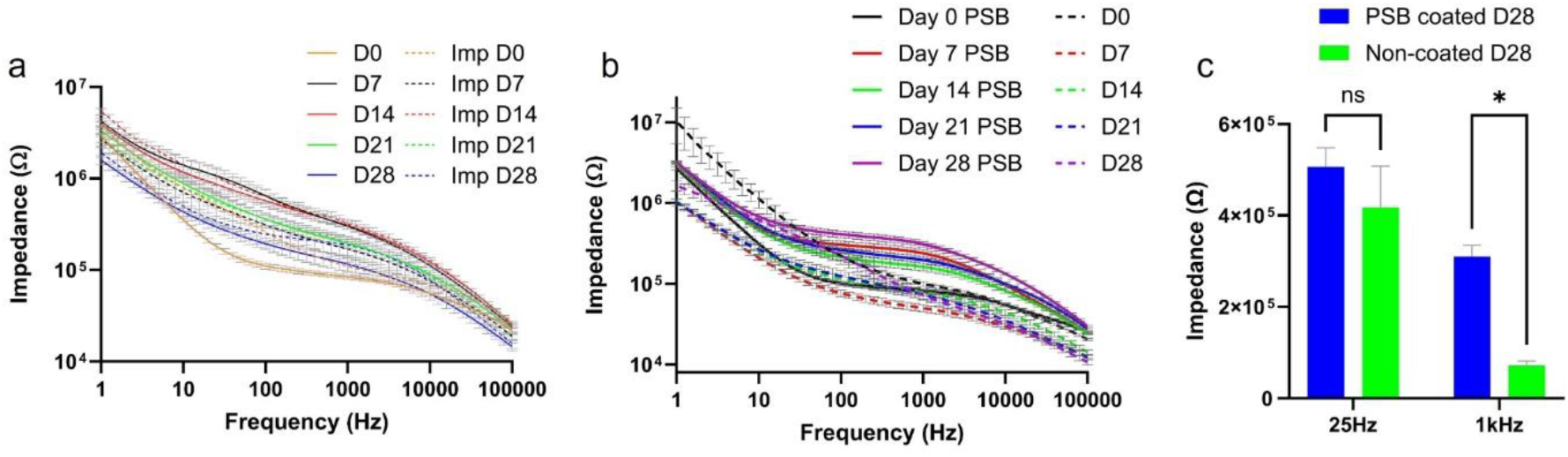
Impedance of PSB PEDOT/CNT coated MEA over 4 weeks. (a), impedance measurement comparison between subcutaneous Ag/AgCl and implanted PSB-coated Ag/AgCl over 4 weeks. (b), Impedance comparison between PEDOT/CNT only and PSB PEDOT/CNT coated MEAs over 4 -weeks. (c) 25Hz and 1kHz impedance comparison at week-4 between PEDOT/CNT only and PSB PEDOT/CNT coated MEAs. [22]

As for the impedance of the PSB PEDOT/CNT MEA, the PEDOT/CNT only MEA had a lower impedance from Day 14 to Day 28 (Figure 6, b). The initial lower impedance of PSB PEDOT/CNT MEA could be the antifouling effects of PSB coating where tissue debris and blood coagulations were prevented from happening (Figure 6, b)[68-70]. The higher impedance of PSB PEDOT/CNT coated MEA could be attributed to better tissue integration where tissue forms more intimate contact with the PSB PEDOT/CNT MEAs than the PEDOT/CNT only MEAs (Figure 6, b)[71, 72].

The 25Hz impedance, frequency of SWV, was previously shown to be negatively correlated with the sensitivity of the sensor [21]. We did not observe a difference between PEDOT/CNT only and PSB PEDOT/CNT group, possibly indicating that the sensor stability is similar between two types of MEAs in vivo. The significantly higher 1kHz impedance of PSB PEDOT/CNT coated MEA could possibly be caused by intimate device/tissue contacts without interference of scar tissue [71, 72].

### Chronic Electrophysiology Recording

To investigate whether PSB coating can provide functional improvements to the flexible MEAs, we quantified and compared peak-to-peak (p2p) amplitude and single unit yield (SUY) between PSB-coated and non-coated groups when animals are under anesthesia (Figure 7). Representative waveforms of isolated single-units over 4-weeks are shown in Figure 8, a. The MEA maintained its ability to record high amplitude units in both cortex and striatum regions over 4 weeks (Figure 7, a), a contrast to our previous finding where recording quality degraded at week-4 in the cortex region [21]. PSB-coated MEAs had significantly higher p2p amplitude than the non-coated ones between Day 7-Day 28 and overall had much higher p2p as well, which also could reinforce the better tissue integration theory (Figure 7, b). The SUY of PSB coated group is significantly better than the non-coated group overall (Figure 7, c). PSB coated group also showed more stable SNR over 4 weeks (Figure S1). These results show that PSB coating can provide better device/tissue integration and enhance the recording performance of implanted MEA chronically.

**Figure 7.**
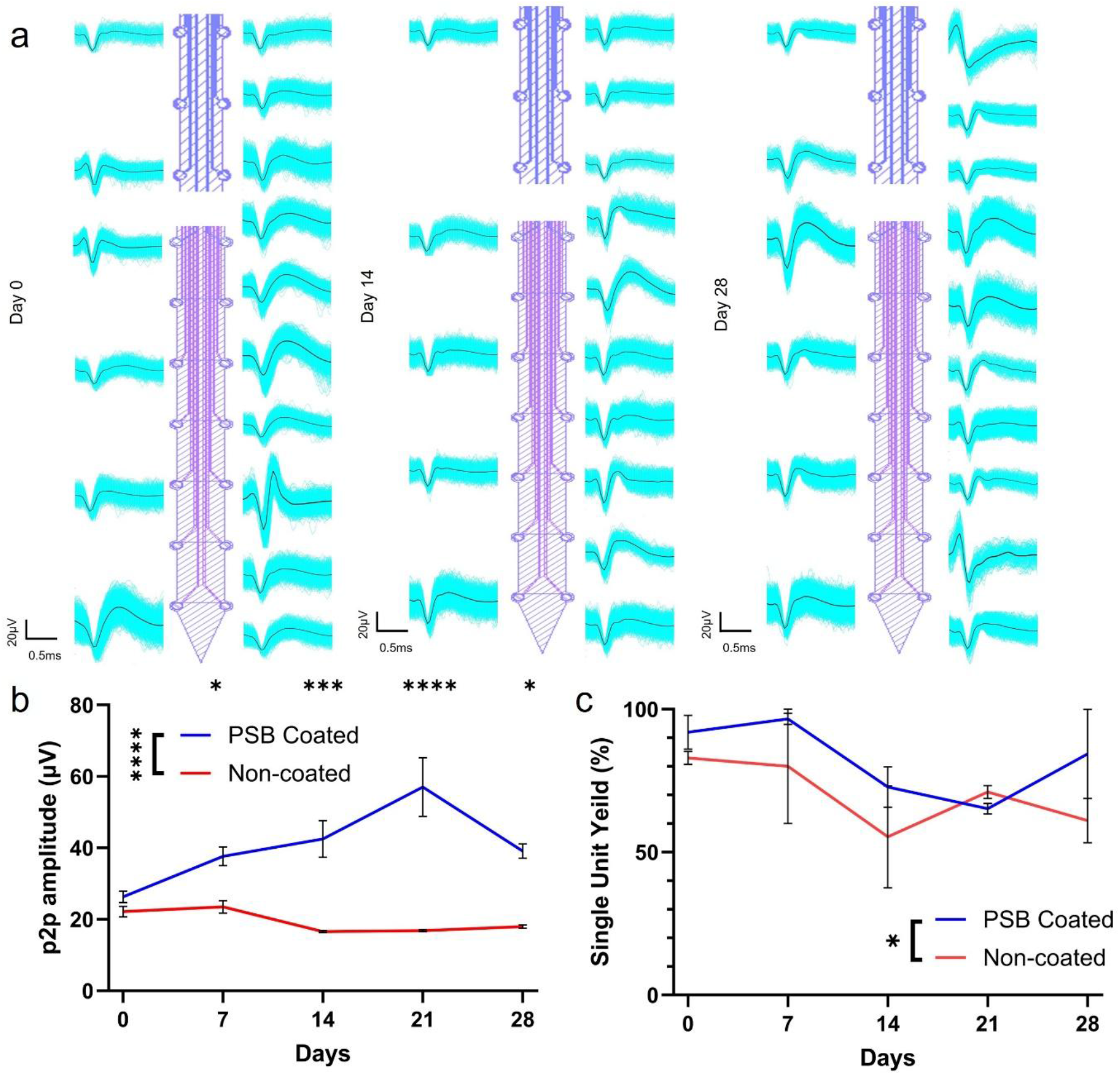
Recording performance of PSB coated MEAs over 4 weeks with animals under anesthesia. (a), Representative waveforms over 4 weeks for PSB-coated MEAs. (b), p2p amplitudes over 4 weeks for PSB-coated and non-coated groups. (Two-way ANOVA, 4 animals in each group, Bonferoni posthoc, * p <0.05, *** p <0.001, **** p <0.0001) (c), SUY of PSB-coated and non-coated groups. (Two-way ANOVA, 4 animals in each group, Bonferoni posthoc, * p <0.05).

**Figure 8.**
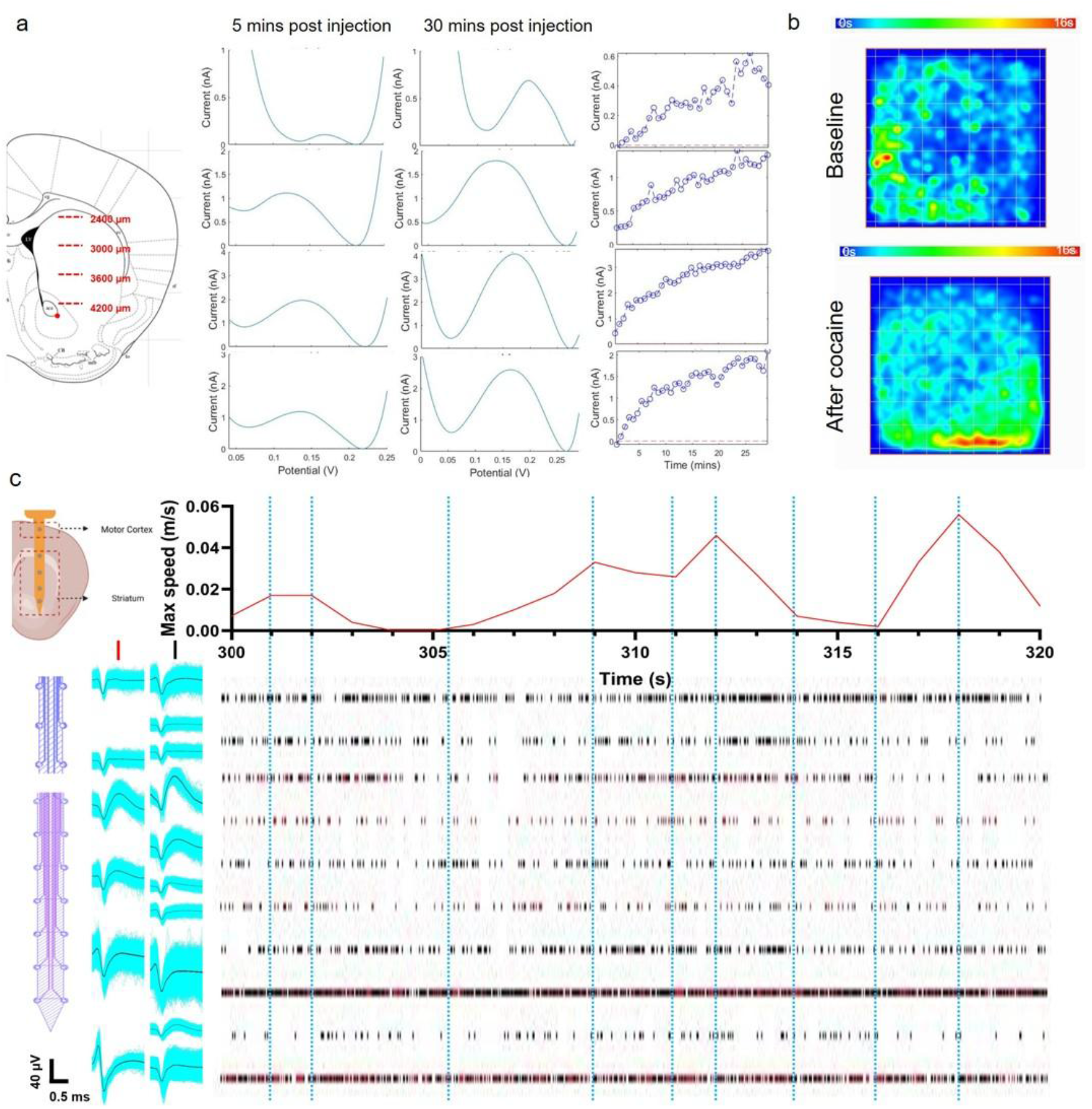
Representative spatial and temporal DA dynamics in a free-moving mouse. The left scheme shows the corresponding tissue depth of the channels shown on the right. (a), 30 mins of DA dynamics after cocaine in a free-moving animal. The SWV waveform is the averaged waveform of first 5 mins and last 5min. (b), Spatial heatmap of animal movements before and after cocaine injection. (c), Representative electrophysiology recording alongside synchronized moving speed in a free-moving animal. Temporarily aligned moving speed and raster plots shows spatially distinct neural activities throughout cortex and striatum. Blue dotted lines are placed to help visualize large movements and corresponding electrophysiological response.

### Electrophysiology and DA Sensing in Free Behaving *Clock*Δ19 Mice

Representative DA sensing and electrophysiology recording in freely moving animals (5 clockΔ19 homozygotes female) are demonstrated in Figure 8. The PSB PEDOT/CNT-coated MEA can stably track DA in free-moving animals for up to 28 days. Spatial and temporal response of DA, average SWV waveforms, and corresponding animal movement tracking on Day 28 is shown in Figure 8, a & b. The cocaine injection resulted in a stably increasing DA concentration across the striatum over 30 minutes (Figure 8, a). The injection also induced elevated motor activities (Figure 8, b). After the cocaine injection, we observe a sustained increase of DA as expected and corresponding increased activity levels from the animal (Figure 8, b). This also helps us to validate that the redox signal we were detecting is indeed DA transients. 20s snippet of the free-moving recording is shown in Figure 8, c, where high-quality single-unit waveforms can be isolated and aligned with animal movements. Variable single units’ activities across the motor cortex and striatum can be observed. The temporal firing patterns of neurons in both cortex regions and the dorsal striatum appear to be entrained with animals’ movements, whereas neurons in the ventral striatum appear to be active regardless of animal motions. These results are consistent with the view that the dorsal-lateral striatum is the motor axis and ventral-medial striatum is the cognitive axis [73-76].

In one clockΔ19 homozygote female animal, we performed continuous DA measurement for 23 hrs to capture the diurnal pattern of DA levels (Figure 9, a). Connected to the same setup described prior in the text, the animal was placed in the behavioral chamber with access to food and water. The animal was allowed to recover from anesthesia and animal movement was tracked during continuous 23hr SWV recording. At the onset of the experiment at ZT6 we observed a peak in DA coinciding with the novelty induced arousal during open field exposure after which DA signals progressively declined to stability throughout the experiment. From approximately ZT8 to ZT10, no animal motion was detected during sleep and DA signal was lower during this period. DA signal stabilized by ZT12 and we observed minor fluctuations in DA signal throughout waking hours until the animal fell asleep at ZT24. Prior to the onset of sleep, we observed a general decline of DA across multiple depths which is consistent with the report that DA concentrations falls following periods of wakefulness (sleep onset) [77] This capability for continuous measurement will be extremely valuable in the study of circadian rhythm, sleep disorders etc.

**Figure 9.**
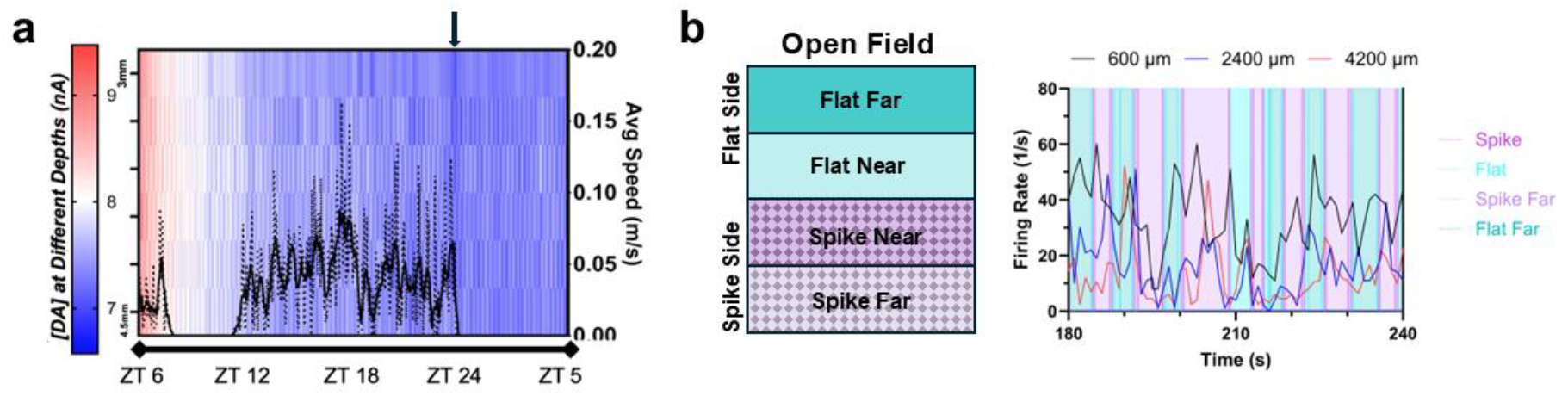
PSB coatings enable stable long term behavioral recordings. (a) Continuous behavior tracking and DA sensing in a ClockΔ19 mouse for 23 consecutive hours. Dashed line: average speed, Solid line: smoothed (10-point avg) average speed. Arrow indicates focal decrease of dopamine prior to sleep onset. B) Neuronal activities during mechanical stimulations on flat vs. spike surfaces. MEA electrodes at depths of 600, 2400, and 4200 μm measure firing rate from single neurons in motor cortex, DS, and NAc, respectively. The testing area is divided using quadrants, where one body length from the area midline shows Spike/Flat crossing. Spike Far and Flat Far are in the far area.

As another proof-of-concept demonstration, a different cohort of 3 clockΔ19 homozygote female animals were implanted with MEAs and we placed one of the implanted mice in a behavioral arena with flat and spiked flooring typically used for mechanical pain evaluation. Fig. 9 b shows single unit firing rates at three different depths of the MEA. We observed changes in firing rate across the motor cortex (600-µm), DS(2400-µm), and NAc(4200-µm) as the animal moved around the arena between the flat and spiked regions. We specifically noticed that the firing rates within the motor cortex are out of phase from those recorded in the DS and NAc as the animal navigates the arena. We also observed that firing activities coincide with spike-to-flat or flat-to-spike crossings.

### Explants and Histology Studies of PSB-coated devices

Explants of both PSB-coated Ag/AgCl wires were imaged using SEM to investigate the coating stability in vivo (Figure 10). The PSB-coating was still present and uniformly distributed over the Ag/AgCl (Figure 10, c), demonstrating excellent stability in vivo. The underlying Ag/AgCl crystal structure was also well-preserved, indicating the PSB coating can also protect the Ag/AgCl reference from brain micromotion-induced physical damage [41]. The explants of PSB-coated MEAs also showed a much cleaner shank surface than the non-coated MEAs with no sign of encapsulated tissue debris (Figure S2).

**Figure 10.**
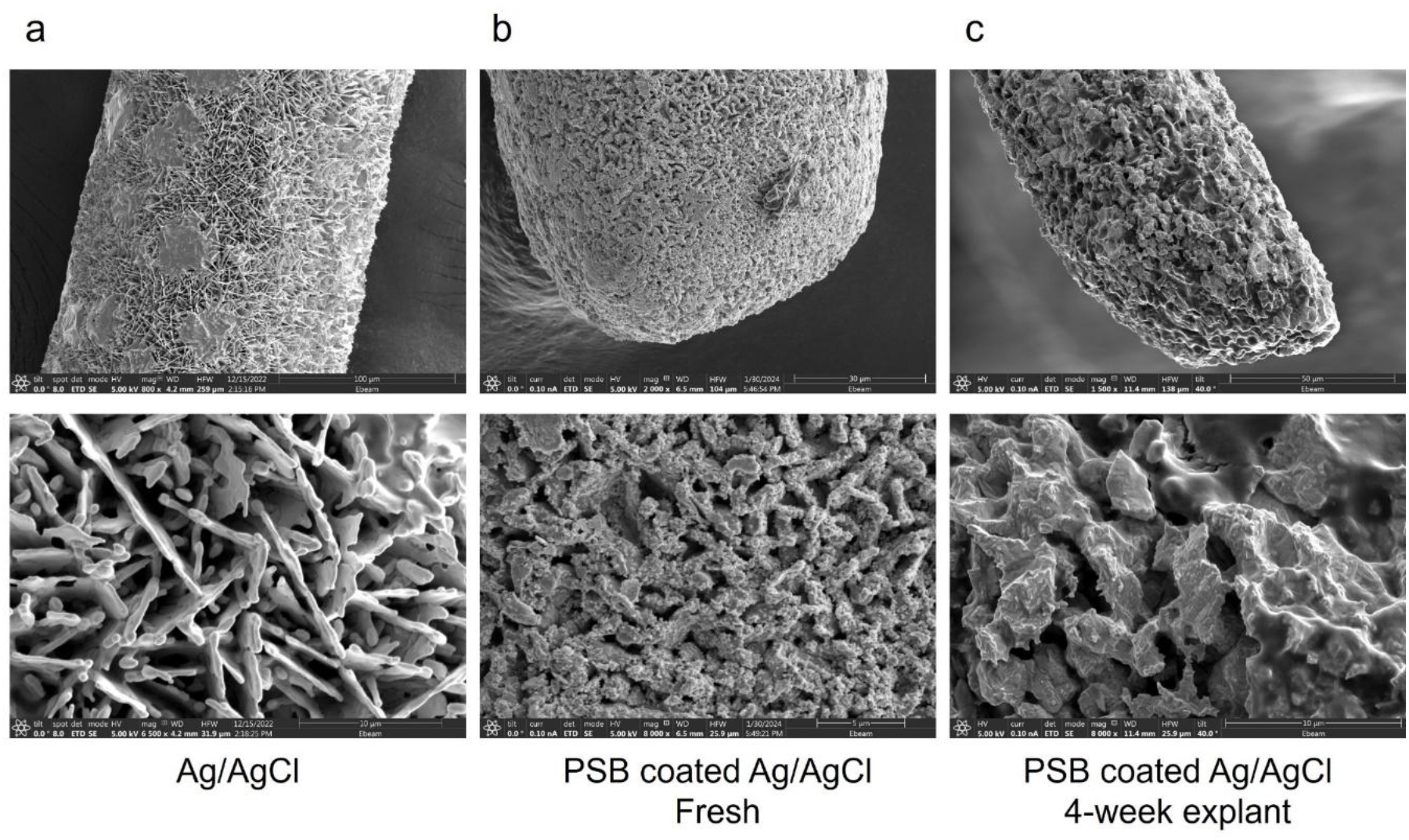
SEM comparison of non-coated Ag/AgCl, freshly PSB-coated Ag/AgCl, and 4-week explanted PSB-coated Ag/AgCl. (a), Ag/AgCl wire, zoomed-in inset shows the crystal structure of AgCl. (b), Fresh PSB-coated Ag/AgCl. The organic-looking PSB polymer layer on top of the AgCl crystal can be easily identified. (c) 4-week explanted PSB coated Ag/AgCl from mice brain. A similar morphology is observed between fresh and 4-week explant. [22]

Additional histological investigation was performed to gauge the surrounding tissue health (Figure 11). Iba-1 (microglia) and GFAP (astrocyte) stains are used to study the inflammatory tissue reactions. Microglia acts as the first responder to brain injury [78], its activity modulates both glial and neuronal tissue responses [79, 80]. Microglia are activated with mechanical and chemical cues that can upregulate proinflammatory cytokines and drive neurons toward excitotoxicity and neurodegeneration [68, 81-84]. The PSB-coated MEAs showed significantly lower Iba-1 intensity level compared to the non-coated group, indicating PSB’s ability to reduce initial microglia reactions that reduce downstream inflammation cascades. The PSB-coated MEAs also has significantly lower GFAP response, indicating decreased astrocyte activity at week-4, indicative of gliosis encapsulating progress of the electrodes that contributes to hindered information flow from and to the devices [68, 83, 85, 86].

**Figure 11.**
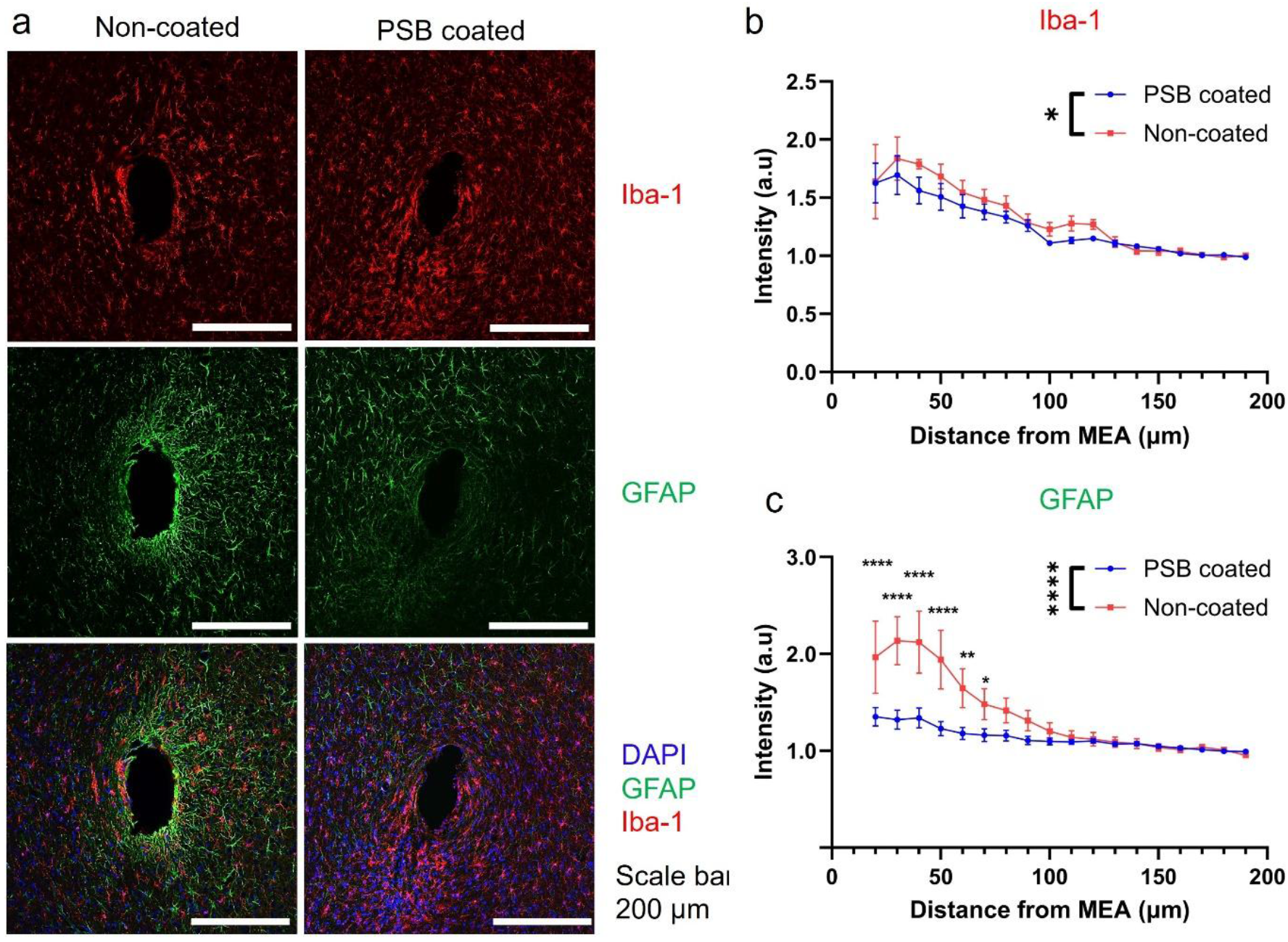
Histology evaluation of tissue inflammation response surrounding the MEA implantation sites. (a), Representative images of Iba-1 (microglia) and GFAP (astrocyte) stained tissue slides. (b). Quantified of Iba-1 intensity up to 200 µm away from the implantation site. PSB coated group showed significantly lower Iba-1 activity than the non-coated group. 10µm bins. Two-Way ANOVA with Tukey’s multiple comparisons test, * p< 0.05. (n = 13, 5 mice for PSB-coated. n = 8, 4 mice for non-coated) (c). Quantification of GFAP intensity up to 200µm away from the implantation site. PSB coated had significantly lower astrocyte reaction compared with non-PSB coated groups. PSB group had significantly lower GFAP intensity than the non-coated groups within 20-80 µm radius. Also, there’s no significant difference in GFAP intensity from 20 µm to 200 µm away from the implantation site for PSB coated group. Whereas non-coated group had significantly elevated GFAP intensity near the implantation site (20-60 µm) compared to far away region (150-200 µm). Significance marker not shown for data clarity. 10µm bins. Two-Way ANOVA with Bonferroni’s multiple comparisons test, **** p<0.0001, ** p<0.005, * p< 0.05. (n = 13, 5 mice for PSB-coated. n = 8, 4 mice for non-coated)

### Conclusion

In conclusion, we demonstrated that PSB coating on Ag/AgCl and MEA surfaces can enable high-quality stable electrophysiology recording and DA sensing in free-moving animals for up to 4 weeks. The PSB coating does not alter the electron transfer kinetics of Ag/AgCl reference and synergistically improves PEDOT/CNT coating quality without affecting sensing towards DA. This technique enables holistic interrogation of DA dynamics, electrophysiological activities, and behavior patterns of free-moving animals, paving the way for a better understanding of the complex interactions between neurocircuit dynamics and behavior generation [22].

## Methods

### Materials

Anhydrous Methanol (MeOH), Diethylammonium diethyldithiocarbamate, Anhydrous Tetrahydrofuran(THF), [2-(methacryloyloxy) ethyl]-dimethyl-(3-sulfopropyl) ammonium hydroxide 97% (SBMA), p-(Chloromethyl)phenyltrimethoxysilane, 3,4-ethylenedioxythiophene (EDOT), α-ketoglutaric acid, Dithiothreitol (DTT), cocaine hydrochloride C-II, nitric acid (95% fuming), and sulfuric acid were purchased from Sigma-Aldrich (St. Louis, MO, USA). Multi-wall carbon nanotubes (CNT) (10-20nm diameter, 10-30µm length, 95%, 200-350 m^2^/g) were purchased from Cheap Tube (Grafton, VT, USA). All materials are used as purchased without further purification.

### Synthesis of N,N-(Diethylamino) dithiocarbamoyl-Benzyl (trimethoxy silane)

SBDC was synthesized using an optimized protocol from previously reported method [63, 65]. In brief, 1.65 g (6mmol) of p-(chloromethyl) phenyltrimethoxysilane (1) and 1.334 g (6 mmol) of diethylammonium diethyldithiocarbamate (2) were dissolved in 10 mL of dry THF separately. Solution of (2) was then slowly added into (1). The solution was stirred for 3 h at room temperature. After filtration and THF evaporation, the product SBDC (3) was obtained as a light-yellow viscous liquid (g, yield). The product is kept cool and in the dark. (1H NMR, (CD3Cl, 40MHz), δ ppm 7.598(2H, d, J = 7.6Hz), 7.419(2H, d, J = 7.6 Hz), 4.558(2H, s), 4.045(4H, d, J = 7.2 Hz), 3.615(9H, s), 1.281(4H, t, J = 6.8 Hz).

### MEA Fabrication

The flexible MEAs were fabricated using the same process from previous work [21, 22]. A 4inch Si wafer with a 500 nm thick SiO_2_ layer (University Wafer Inc., USA) was first cleaned by sonicating in acetone, isopropanol, and DI water sequentially for 5 mins, respectively. 1). The cleaned wafer was dried on a hot plate at 95°C for 3 mins, then the surface was cleaned and activated by O_2_ plasma using a reactive ion etcher (RIE, Trion Phantom III LT, Clearwater, FL, USA) for 90s at 200mTorr pressure and 150Watts power. The wafer was then spin-coated with polyimide (PI) HD4100 (HD MicroSystems L.L.C. NJ, USA) at 3000rpm for 1min and soft baked at 95°C for 1 min to evaporate solvent, then the wafer was exposed with a customized mask using a Quintel Q4000 mask aligner (NEUTRONIX QUINTEL, CA, USA) at a dose of 400mJ/cm^2^. After exposure, the PI layer was post-baked at 95°C for 2 mins, developed using PA-401D developer (HD MicroSystems L.L.C. NJ, USA) for 1min. The wafer was then rinsed and cleaned with PA-400R (HD MicroSystems L.L.C. NJ, USA) and dried and baked at 95°C for 1min. The wafer was cured at 200°C for 30 mins and 350°C for 1 hr in a OTF 1200 Series tube furnace purged with N_2_ (MTI Corporation, CA, USA). 2). The wafer was then treated with O_2_ plasma to clean, activate, and roughen the PI with RIE for 60s at a pressure of 200mTorr and 150 W power. The treated wafer was then spin-coated with AZ P4210 photoresist (MicroChemicals, Germany) at 4000 rpm for 1min and baked at 105°C for 2 mins for resist curing. After baking, the wafer was exposed using MLA 100 (Heidelberg Instruments, Germany) with a dose of 250 mJ/cm^2^, then developed using AZ400k 1:4 developer (MicroChemicals, Germany), cleaned by water rinse, and dried with N_2_ gas flow. A mild 120 s RIE O_2_ plasma treatment at a pressure of 600 mTorr and 60 W power was performed to clean the surface before metal deposition. A 15nm Ti adhesion layer, 100 nm Au, and 20nm Pt layer were evaporated on the wafer using an Electron Beam Evaporator Plassys MEB550S (Marolles-en-Hurepoix, France). The metal was then lifted off in acetone overnight. 3). The next day, the wafer was first rinsed with water, dried under N_2_ flow, and cleaned by O_2_ plasma for 60 s at 200 mTorr and 150 W, then spin-coated with HD4100 for the middle PI insulation layer at 5000 rpm for 1 min and soft baked at 95°C for 5 min. The wafer was then flood exposed using a mask aligner with a dose of 400 mJ/cm^2^, post-baked, and cured at 200°C for 30 mins and 350°C for 1hr in a tube furnace purged with N_2_. 4). Then step-2 was repeated to deposit another layer of metal traces and sites. 5). The wafer was spin-coated with HD4100 for the last PI insulation layer at 3000 rpm for 1 min and soft baked at 95°C for 5 mins. The wafer was then flood exposed using a mask aligner with a dose of 400 mJ/cm^2^, post-baked, and cured at 200°C for 30mins and 350°C for 1hr in a tube furnace purged with N_2_. 6). The wafer was then spin-coated with AZ P4620 photoresist at 2000 rpm for 1min and baked at 105°C for 10 mins for resist curing. After baking, the wafer was exposed using MLA with a dose of 950 mJ/cm^2^, then developed using AZ400k 1:4 developer, cleaned by water rinse, and dried with N2 gas flow. The sites and contact pads are dry etched open using RIE (200 mTorr, 180 W, 180 s, 50 sccm O_2_, 2 sccm SF_4_. 180 s etching with 120 s cool down each cycle, 5 cycles). 7). The flexible PI MEAs were released from the wafer using buffered oxide etchant (1:7) (MicroChemicals, Germany) in an acid hood for 8hr to etch away the SiO_2_ layer. Customized PCBs (Wavemed, Italy) were mounted with one 23-contact ZIF (Digikey, USA) and two 16-channel Omnetics connectors (Omnetics connector corporation, MN, USA). ZIF connectors were used to interface with flexible MEAs, and Omnetics connectors were used to interface with the potentiostat and electrophysiology recording system.

### Surface Modification and PSB Grafting

Flexible PI probes were first rinsed with anhydrous methanol three times and air dried for 15 min. After which, all samples were exposed to O_2_ plasma for 10 mins and immediately immersed in 4 mL of SBDC (2 mg·mL^-1^, 5 × 10^-3^ M) anhydrous MeOH solution, sealed and covered with aluminum foil, and left overnight. The next day, probes/wafers were rinsed with anhydrous MeOH three times, kept in a petri-dish, air dried, and covered with foil before use. 4 mL of PBS solution was prepared in a quartz tube, sealed, and degassed with N_2_ for 30 mins. Meanwhile, 1 g of SBMA monomer was prepared in a test tube, and the powder was vacuumed for 15 min, then purged with N_2_ gas, repeated three times. The degassed PBS solution was then transferred with a syringe and needle into an SBMA-containing tube, then well-mixed SBMA PBS solution was transferred back to a quartz tube. Probes were then lowered into SBMA solution under N_2_ flow. The apparatus was then sealed and placed under the UV lamp (100 w, standard filter (320–500 nm), series 1000 Omni Cure) for 1 h under 100% power output setting. The distance between the light source and the substrate is 1.5 cm and the area covered by light is 63.17cm^2^. For probe coating, the quartz tube rotated 180° every half hour to ensure uniform polymer growth on the front and back. After polymerization, probes were rinsed with DI water, dried under N_2_, covered with aluminum foil and stored at room temperature before coating with PEDOT/CNT [22].

For Ag/AgCl PSB coating, 4 g SBMA monomers were dissolved in 20 mL 1 x PBS solution. 70 mg of Dithiothreitol (DTT) and 10 mg α-ketoglutaric acid were dissolved into the same solution. The solution was then stirred and partially polymerized under a UV lamp (100 w, standard filter (320–500 nm), series 1000 Omni Cure) for 1 h under 100% power output setting (1.5 cm away from lamp). Ag/AgCl wires were then manually dipped into the partial polymerized solution and exposed to the same UV lamp for 30 s. This process is repeated 3 times. Then the samples are transferred to a UV oven (Form Wash, Boston, MA, US), cured for 3hrs at 70°C. The PSB-coated Ag/AgCl wires were rinsed with sterile PBS and soaked in 70% ethanol for 10 mins before being used for surgery [22].

### Electrochemistry Methods

All electrochemical procedures were conducted using a three-electrode design (working electrode: individual MEA electrodes; reference electrode: Ag/AgCl; counter electrode: Pt (*in vitro*)/stainless-steel bone screw (*in vivo*). An Autolab potentiostat/galvanostat, PGSTAT128N (Metrohm, Herisau, Switzerland) was used for all square wave voltammetry (SWV) procedures and electrochemical Impedance spectroscopy (EIS) measurements (*in vivo* and *in vitro*). SWV potential was swept from −0.2v to 0.3V using a 25 Hz pulse frequency, 50 mV pulse amplitude, and a 5mV step height. A liner scan from 0.3V to 0V at 1v/s was applied after SWV waveform. Potential was held at 0V between scans. *In vitro* DA calibrations were performed using freshly prepared DA standard solutions dissolved in 1x PBS (50 nM, 100 nM, 150 nM, 250 nM, 500 nM).

### CNT Functionalization and PEDOT/CNT Deposition

CNTs were functionalized following previous methods [87, 88]. In brief, 200 mg of multiwalled carbon nanotubes was added to 25 ml of concentrated nitric acid and 75-ml concentrated sulfuric acid. This solution was sonicated for 2 h, and then stirred overnight at 35 °C. The solution was dialyzed in a DI water bath until the solution became pH neutral. The water bath was changed every 12 hours. Samples were vacuum dried and stored at 4 °C.

For PEDOT/CNT deposition, 1 mg/mL of functionalized CNTs was resuspended in DI H2O by sonication for 10 min. EDOT was added to this solution to a concentration of 0.01 M. The solution was then sonicated for 10 minutes using a Q500 probe sonicator (Qsonica L.L.C, Newtown, CT, USA). Electrochemical deposition was performed using chronocoulometry. The applied voltage is 0.9V, with a charge cut off at 150mC/cm^2^.

### Scanning Electron Microscope (SEM) Characterization

SEM images were obtained with an FEI Scios Dual Beam System (ThermoFisher Scientific, Waltham, MA, USA) with a 5 kV, 0.1 nA beam and a 7 mm working distance.

### Animal Housing and Breeding

Mice were group housed on a 12/12 light/dark cycle (lights on 7 a.m., lights off 7 p.m.) with food and water ad libitum. *Clock*D19 mice on a Balb/c mixed background were bred as heterozygotes to produce WT and homozygous MU littermates. Female *Clock* mutant (*Clk*Δ19/*Clk*Δ19), 6-10 weeks old, were used in all studies.

### Surgical Procedure

All animal work was performed under the guidelines of the University of Pittsburgh Institutional Animal Care and Use Committee (IACUC). The approved protocol ID is 22109970. Mice were anesthetized under isoflurane (2.5%) and head-fixed in a stereotaxic frame (David Kopf Instruments, Tujunga, CA, USA). Animal body temperature was maintained at 37 °C using an isothermal pad connected to a SomnoSuite system (Kent Scientific Corporation, Torrington, CT, USA). Heart rates were monitored using the SomnoSuite system as well. A holder was used to secure the custom-designed PCBs. The TDT recording system (RX5, 16-channel Medusa amplifier, Tucker Davis Technologies (TDT), Alachua, FL) was used to record electrophysiology data. Three skull screws were carefully positioned above the left striatum, right visual cortex, and left visual cortex of the mice as illustrated in Figure 4, a). A 0.7mm diameter window above the ventral striatum of the right hemisphere was opened using a motorized drill. The coordinates for the center of the window were 1 mm posterior to Bregma and 1.1 mm lateral to the midline. The flexible shank was temporarily glued to a 50µm diameter tungsten wire shuttle with 30% polyethylene glycol (PEG, MW 20KDa). The PCB/MEA assembly, tungsten wires, and PEG solution were UV sterilized for 10 minutes before surgery, and tungsten wires were attached to MEA shanks using sterilized PEG before implantation. The fully assembled device was inserted into the brain using a motorized micromanipulator from NeuralGlider (Actuated Medicine, Inc, Bellefonte, PA, USA), 4.5mm deep into the brain. A Pt counter wire was connected to the skull screw on the contralateral hemisphere for electrophysiology recording. Another Pt wire was connected to the screw above the contralateral visual cortex. Both wires were soldered to the corresponding ground wires on the PCB [22].

### *In Vivo* DA Sensing and Electrophysiology Recording

Animals were anesthetized with 2% ISO, then the head-stage for either potentiostat or recording system was connected to the animal head through and imbedded PCB using Omnetics connectors. For each session, EIS was first measured with a fresh Ag/AgCl inserted subcutaneously. Electrophysiology recording was collected for a 5min duration. EIS was measured again using an implanted subcutaneous Ag/AgCl reference. Animals were then put into an open field and allowed to wake up. A 20mins SWV DA detection was performed after the animal had fully recovered from anesthesia. After free-moving DA sensing, the system was switched to electrophysiology recording and 5 mins of free-moving electrophysiology data were recorded. A cocaine injection (10mg/kg) was given on day 28 after the free-moving electrophysiology session. Cocaine was dissolved in PBS and delivered through intraperitoneal injection. Additional 30 mins free-moving SWV measurement was performed after cocaine injection. The neural signals from recording microelectrodes were amplified using a 16-channel Medusa preamplifier and recorded with an RX5 processor at 25kHz [22].

### Immunohistochemistry

According to the University of Pittsburgh IACUC-approved methods, mice were sacrificed at the terminal timepoint (4 weeks). Each animal was deeply anesthetized using an 80-100 mg/kg ketamine, 5-10 mg/kg xylazine cocktail. Once mice were unresponsive to tail/toe pinches, animals were transcardially perfused using phosphate-buffered saline (PBS) flush at <100 mmHg followed by 4% paraformaldehyde (PFA) at <100 mmHg. Mice were decapitated and the skulls were removed to post-fix the brain in a 4% PFA at room temperature for 12hr. Then, brains were cryoprotected via sucrose gradient dehydration via soaking in a 15% sucrose (Sigma-Aldrich Corp., St. Louis, Missouri) bath at 4 °C overnight, followed by a 30% sucrose solution for 24 h. Brains were then carefully frozen in a 2:1 20% sucrose in PBS:optimal cutting temperature compound (Tissue—Plus O.C.T. Compound, Fisher HealthCare, Houston, TX) blocking media blend with dry ice chilled 2-methylbutane. Frozen tissue was then transversely sectioned at a thickness of 25 μm spanning the depth of implanted probes using a cryostat (Leica CM1950, Buffalo Grove, IL).

Cortical sections of implanted and non-implanted hemisphere were mounted on the same slide for comparison and staining for each antibody combination was performed at the same time to minimize variability. Antibodies to visualize astrocytes (GFAP, 1:500, Z033401 Dako), macrophage/microglia IBA-1(microglia, 1:500, NC9288364 Fisher).

Tissue sections were rehydrated in 1 x PBS for 2×5 min. The tissue was then incubated in 0.01 M sodium citrate buffer for 30 min at 60 ° C. Then, a peroxidase block (PBS with 10% v/v methanol and 3% v/v hydrogen peroxide) was performed for 20 min at room temperature (RT) on a table shaker. Next, tissue sections were incubated in carrier solution (1 X PBS, 5% normal goat serum, 0.1% Triton X-100) for 30 min at RT. Lastly, the tissue sections were blocked with Alexa Flour 647-conjugated AffiniPure Fab Fragment goat anti-mouse IgG (IgG, 1:16, 115-607-003 Jackson ImmunoResearch Laboratories, Inc.) or Fab fragment only (1:13, 115-007-003, Jackson ImmunoResearch Laboratories, Inc.) for 2 hours then rinsed 6 times each 4 minutes. Following blocking, sections incubated in a primary antibody solution consisting of carrier solution and antibodies listed above overnight (12-18 hr) at RT. Sections were then washed with 1 x PBS for 3 × 5 min and incubated in carrier solution and secondary antibodies (1:500, Alexa Flour 488 goat-anti mouse, Invitrogen, and 1:500 Alexa Flour 568 goat-anti rabbit, Invitrogen, 1:500 Alexa Flour 633 goat-anti chicken, Invitrogen) for 2 hours at RT. Then sections were rinsed with PBS for 3 × 5 min and exposed to Hoechst (1:1000, 33342 Invitrogen) for 10 min and washed in PBS for 3 × 5 min, then covered by coverslip with Fluoromount-G (Southern Biotech, Associate Birmingham, AL).

### Data Analysis

Each SWV response was first filtered using a zero-phase, forward and reverse (using the filtfilt function on MATLAB), low-pass, third-order Butterworth digital filter with the 3 dB cutoff at a normalized frequency of 0.2 (2 Hz). The fit for the linear baseline was determined using a two-step peak extraction method consisting of an iterative peak localization algorithm. First, a linear baseline was initialized with two signal points on either side of a user-selected peak maximum voltage (∼0.18 V). Signal points used to construct the baseline were iteratively updated to produce a final baseline which maximized the subtracted peak amplitude. The resulting linear fit intersected boundary points at either side of the DA redox peak profile. The five data points immediately adjacent to the upper and lower bounds were then modeled using linear fitting and subtracted from the raw SWV response for the purpose of peak extraction. All extracted peak currents were converted to DA concentration using in-vitro calibration curves.

Raw neural recording data was filtered between 300 Hz to 10 k Hz. Threshold crossing events were identified by using a fixed negative threshold value of 3.5 standard deviations. Plexon offline sorter (Plexon Inc Dallas, TX, USA) was used to identify single units. A 3D PCA feature space was used to identify waveform features and K-means clustering method was used to identify individual units. K-means was set up using an adaptive standard EM between 2 to 5. The Signal-to-Noise (SNR) ratio of sorted units were calculated and units with a SNR above 4 were included in analysis. The SNR is calculated using the equation below:

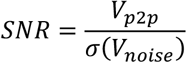

where V_p2p_ is peak to peak amplitude of single units, and σ(V_noise_) is the standard deviation of the high pass filtered stream.

The single unit yield (SUY) is calculated using the equation below:

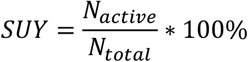

Where N_active_ is number of electrode sites that recorded single units, N_total_ is the total number of functional electrode sites.

A customized MATLAB script was used to calculate SNR, SUY, spike rates, and p2p amplitudes. Statistical analysis and result plotting were done using GraphPad Prism 10.0.0 and MATLAB 2019a. Data described in-text are mean ± SM unless specified otherwise. Schemes were drawn with BioRender.

## Supplementary Materials

Supporting Information is available from the Wiley Online Library or from the author.

## Author Contributions

Conceptualization: B.W, and X.T.C.; methodology: B.W and T.D.; validation: B.W; formal analysis: B.W; investigation: B.W, C.M and X.T.C; resources: Y.X., C.M and X.T.C; data curation: B.W, and C.T. writing—original draft preparation: B.W and C.T.; writing—review and editing: B.W, C.T, Y. X, C.M, and X.T.C.; supervision: C.M and X.T.C; project administration: B.W., C.M, and X.T.C; funding acquisition: Y.X., C,M and X.T.C. All authors have read and agreed to the published version of the manuscript.

## Funding

This work was supported by the National Institutes of Health BRAIN R01NS110564, R21DA049592, R21NS123937 to Dr. X. Tracy Cui and R01MH106460, R01DA039865 to Dr.

Colleen McClung. Dr. Thompson was supported by an NIH Ruth L. Kirschstein National Service Award (T32GM075770, PI/PD: Yan Xu).

## Data Availability Statement

The data presented in this study are available on request from the corresponding author.

## Acknowledgments

We would like to thank the staff at the Nanoscale Fabrication and Characterization Facility of the University of Pittsburgh for their technical support for MEA fabrication and SEM instruments. We would like to thank the staff at the Center for Biologic Imaging of the University of Pittsburgh for their technical support for confocal microscopes.

## Conflicts of Interest

The authors declare that they have no known competing financial interests or personal relationships that could have appeared to influence the work reported in this paper.

## Notes

### Competing Interest Statement

The authors have declared no competing interest.

### Summary of Updates

Correcting formatting errors and figure numbering errors. Added additional references.

