## supporting info for "Zwitterionic polymer coating enabled chronic dopamine sensing and electrophysiology recording in free-moving mice" (from another version of this preprint: version 1, 2026-02-10)

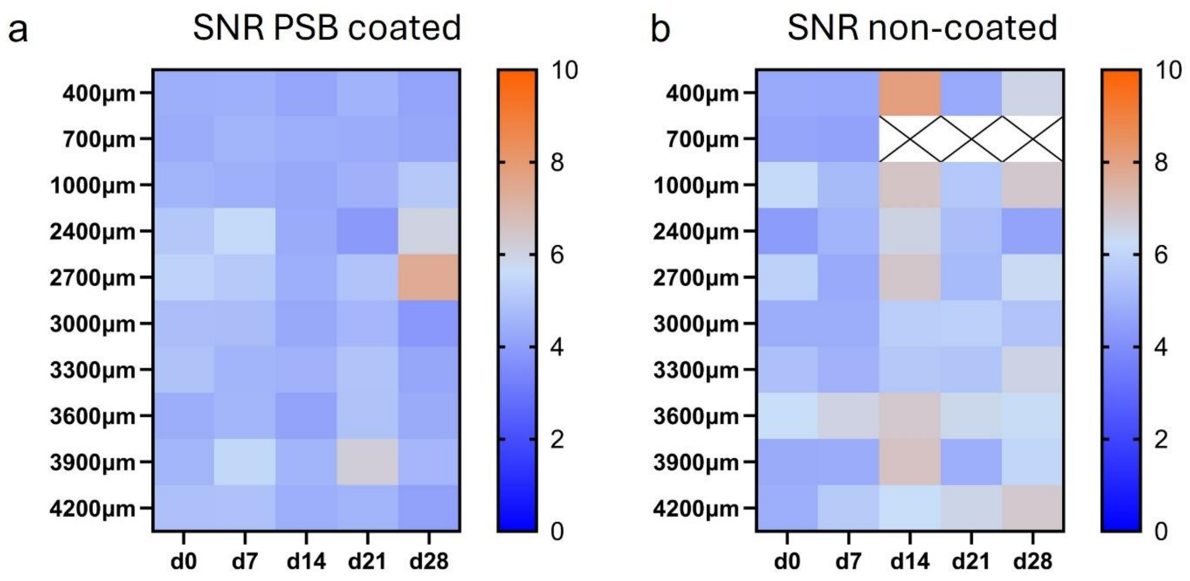


Figure S1. Spatial and temporal patterns of SNR for both groups. (a), PSB coated group showed a much more stable SNR over 4 weeks. (b), Non-coated group showed varying SNR and even loss of channels after 2 weeks.

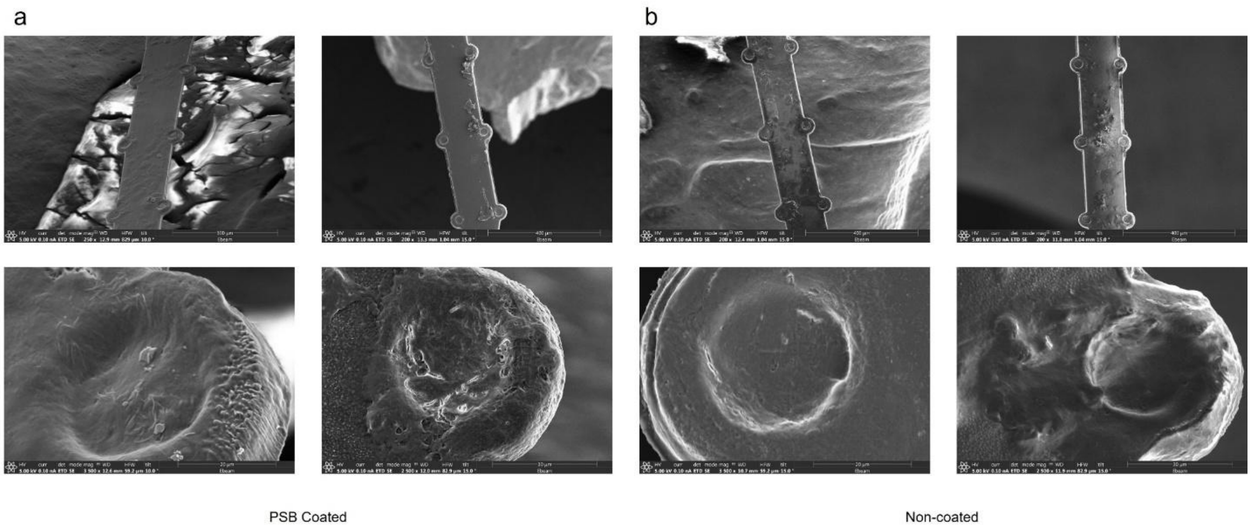


Figure S2: SEM comparison of PSB coated and non-coated explants. (a) PSB coated, 4-week explants. (b) non-coated, 4-week explants.
